# Structural and functional characterization of oligomeric states of proteins in RecFOR pathway

**DOI:** 10.1101/2019.12.17.877662

**Authors:** Santosh Kumar Chaudhary, Mohanapriya Elayappan, Jeyaraman Jeyakanthan, Kanagaraj Sekar

**Affiliations:** Department of Physics, Indian Institute of Science, Bangalore 560012, India; Department of Computational and Data Sciences, Indian Institute of Science, Bangalore 560012, India; Department of Bioinformatics, Alagappa University, Karaikudi 630003, India

**Keywords:** Dimeric TtRecF, Tetrameric TtRecR, Binding affinity, Heterohexamer, RecOR complex, RecFR complex

## Abstract

RecFOR pathway is the principal repair pathway for double strand break and single strand gap repair in *Thermus thermophilus*. RecF and RecR exist as monomer and dimer in solution, interestingly; they undergo condition-dependent dimerization and tetramerization, respectively during the DNA break repair. However, their importance in protein-protein and protein-DNA interactions remains elusive. In this study, the three-dimensional crystal structures of the wild type RecF and RecR proteins are determined. Thereafter, the structural information is used to mutate the interface residues to cysteine to stabilize the dimeric and tetrameric states of the RecF and RecR proteins, respectively. A comparative study for their interactions with other cognate proteins and ssDNA in native and SSB (single strand binding protein) bound states was performed. RecF or RecFR complex displays a negligible affinity towards ssDNA. Conversely, the RecF mutants and its complexes with wild type RecR showed affinity towards ssDNA, suggesting, distinct modes of interaction of RecF and RecFR complex for ssDNA binding. In the presence of RecO, the stabilized tetrameric RecR showed a lower binding affinity for ssDNA as compared to the SSB bound ssDNA, indicating the importance of tetrameric RecR in stabilizing the RecOR complex on the SSB coated ssDNA. This provides an insight into the reduction of the binding affinity of SSB proteins with the ssDNA, which in turn enhances the recruitment of RecA for strand exchange.

## Introduction

The RecFOR pathway proteins play an important role in the restart of stalled replication fork and homologous recombination in prokaryotes (Bidnenko *et al*., 1999). RecBCD and RecFOR are the two major repair pathways present in prokaryotes. The RecBCD pathway is mainly responsible for the double strand DNA (dsDNA) break repair, whereas, the RecFOR pathway is required for single strand DNA (ssDNA) gap repair. However, in the absence of RecBCD pathway proteins, the RecFOR pathway also participates in dsDNA break repair. In contrast to other prokaryotes, the RecBCD pathway is absent and RecFOR is the principal DNA repair pathway in *D. radiodurans*, *B. subtilis* and *T. thermophilus*. In a DNA break event, the helicase RecQ unwinds the DNA (Harmon & Kowalczykowski, 1998) and the RecJ protein creates a 3’ overhang by 5’-3’ exonuclease activity (Lovett & Kolodner, 1989). The 3’ overhang of DNA is subsequently coated with the SSB (single strand binding) proteins. The RecF, RecO and RecR proteins facilitate the loading of the RecA protein on SSB coated 3’ overhang (Morimatsu & Kowalczykowski, 2003). The RecA protein removes SSB proteins from the 3’ tail of the DNA and forms a nucleoprotein filament. The nucleoprotein filament mediates heteroduplex formation between the ssDNA and the complementary homologous dsDNA for homologous recombination (Shibata *et al*., 1979; Handa *et al*., 2009; McEntee *et al*., 1979).

The RecR protein from the RecFOR pathway is reported as an essential repair mediator protein for all the homologous recombination and DNA repair pathways in *M. tuberculosis* (Gupta *et al*., 2015). RecR exists as a dimer in solution; however, the tetrameric assembly of RecR is reported in the crystal structures from *D. radiodurans* and *T. tengcongensis* (Tang *et al*., 2012; Lee *et al*., 2004). The tetrameric assembly of RecR monomers generates a central cavity, large enough to pass a dsDNA. The tetrameric assembly of RecR is required for the interaction with RecF and RecO proteins (Honda *et al*., 2008; Timmins *et al*., 2007). The proteins, RecF and RecO interact with the RecR in the ratio 4:2 (4RecR:2RecO/2RecF) (Tang *et al*., 2012; Timmins *et al*., 2007). RecF and RecO proteins exist as a monomer in solution. The RecF protein shows ATP dependent dimerization and dimer dependent ATPase activity upon DNA binding (Koroleva *et al*., 2007). The monomeric RecF protein does not show any ATPase activity and is active only in the presence of DNA (Tang *et al*., 2018). A recent study on RecF from *T. tengcongensis* revealed that the RecF forms a dimer in the presence of ATP, whereas in the presence of non-hydrolysable ATP analogue (ATPγS), the dimeric interface changes and the helical region of DNA binding domain is involved in dimerization (Timmins *et al*., 2007).

RecFOR pathway proteins form various conditional homo and heteromeric complexes in solution. The roles of their oligomerization in protein-protein and protein-DNA interactions are not clear. In this study, the RecF and RecR proteins from *T. thermophilus* (TtRecF and TtRecR) are successfully crystallized and the high-resolution crystal structures of TtRecF and TtRecR proteins are solved. A dimeric interface of TtRecF is modeled based on the dimeric crystal structure of RecF from *T. tengcongensis*. To stabilize the dimeric state of TtRecF protein, cysteine mutations are performed on its interface residues. Similarly, the tetrameric TtRecR structure is also stabilized by di-sulfide bridges. The interaction properties of the cysteine mutants of RecF and RecR have been compared with the respective wild type proteins. The stabilized tetrameric RecR shows distinct binding properties with the ssDNA, when complexed with RecF and RecO proteins as compared to the wild type RecR. Wild type RecF shows negligible affinity towards ssDNA, interestingly, the mutant RecF shows affinity with the ssDNA and it increases in the presence of the wild type RecR. This suggests that RecF might have distinct binding modes for the ssDNA and dsDNA. Overall, the present study provides insights into the roles of the oligomeric states of RecF and RecR proteins in protein-protein and protein-DNA interactions.

## Materials and methods

### Mutagenesis, Protein expression and purification

The *rec*F, *rec*O, *rec*R and *ssb* genes from *T. thermophilus* cloned in pET11a vector were brought from RIKEN, Japan. The mutants TtRecR_G169C_, TtRecF_D57C_ and TtRecF_S261C_ were generated using the inverse PCR technique. The primer details are provided in supplementary Table S1. The proteins were expressed in *E. coli* BL21DE3 (RIL) cells. These cells were harvested and suspended in lysis buffer. The lysis buffer used for TtRecR, TtRecR_G169C_, TtRecO and TtSSB consists of Tris-HCl pH 7.5, 150 mM NaCl and 5mM β-merchaptoethanol, whereas RecF, RecF_D57C_ and RecF_S261C_lysis buffer consists of Tris-HCl pH 7.5, 500 mM KCl, 25 mM Sucrose, 5 mM lysozyme and 5 mM β-merchaptoethanol. TtRecR, TtRecR_G169C_ and TtSSB were purified using anion exchange column and TtRecF, TtRecF_D57C_, TtRecF_S261C_ and TtRecO were purified using HiTrap Heparin HP column (GE healthcare). All the purified proteins were further polished using gel filtration column (Hi-load 16/600 Superdex200pg, GE healthcare) pre-equilibrated with the buffer containing 20mM HEPES pH 7.5 and 150 mM NaCl (Supplementary Fig. S1). The protein concentrations were measured using Bradford assay.

### Crystallization, Data Collection and processing

The proteins TtRecF and TtRecO were successfully crystallized using under oil microbatch (greiner plate) technique. The drops contained 2 µl of protein and 2 µl of the crystallization condition. Silicon and paraffin oil (3:1) of 7ml was used in the plate. The crystallization condition of TtRecF consists of 0.1M Sodium malonate pH 7.0, 0.1M HEPES pH 7.0 and 0.5 % v/v Jeffamine ED-2001 pH 7.0 and TtRecR in 0.2M Potassium chloride, 0.05 M HEPES pH 7.5 and 35% v/v Pentaerythritol propoxylate. The data was collected at home source (Molecular Biophysics Unit, Indian Institute of Science, Bangalore, India) and BM14 beam line (European Synchrotron Radiation Facility (ESRF), Grenoble, France). The data sets were processed using iMOSFLM (Xu & Freitas, 2009), xia2 (Holm & Laakso, 2016), DENZO and SCALEPACK (Tang *et al*., 2018).

### Structure solution and refinement

The three dimensional crystal structures of DrRecF (PDB id – 2O5V) and DrRecR (PDB id – 1VDD) were used as starting models for molecular replacement of TtRecF and TtRecR, respectively, using the program PHASER (Moncalian *et al*., 2004). The sequence identities between DrRecF and TtRecF, and DrRecR and TtRecR were 40 % and 50 %, respectively. The crystal lattice of the wild type TtRecF and TtRecR belong to the primitive orthorhombic space groups, P22_1_2_1_ and I222, respectively, with one molecule in the asymmetric unit. The reciprocal space refinement was carried out using maximum likelihood method implemented in REFMAC program (Koroleva *et al*., 2007), integrated with CCP4 program suite (Lee *et al*., 2013; Baker *et al*., 2001), while real space refinement was performed using COOT (Lee *et al*., 2013). The model was submitted to PDB_REDO (Tang *et al*., 2018; Webb *et al*., 1999) for geometry optimization followed by cycles of refinement using REFMAC and COOT. A total of 5 % of the reflections were set aside for R-free calculations.

### Native PAGE and Electro Mobility Shift Assay (EMSA)

TtRecR (10 µM), TtRecO (10, 20, 40 µM) and TtRecF (10, 20, 40 µM) were incubated in 25 µl of 20 mM HEPES (pH 7.5) and 150mM NaCl. After 10 min at 50°C, the samples were analyzed by polyacrylamide gel electrophoresis (PAGE) using 8 % gel under non-denaturing conditions. The proteins were visualized by Coomassie Brilliant Blue R-250 (CBB) staining. Fluorescein (FAM) labelled dT_35_ at the 5’ and 3’ of the oligonucleotide was used for the EMSA experiments. 2 µM of FAM labelled dT_35_ was used for the experiments. Similar buffer and protein concentrations were taken, as used for native-PAGE analysis. DNA was visualized by Bio-rad chemidoc using fluorescein filter.

### Bio-layer Interferometry

ForteBio’s Super Streptavidin biosensors were used for the immobilization 50 nM of biotinylated ssDNA (Biotin-5’GCAATGAAGTACGCTTGCCAGCTGCAGTCATGATTG3’) (Sultana & Lee, 2015). The binding affinities of TtRecR, TtRecR_G169C_, TtRecF, TtRecF_D57C_, TtRecF_S261C_, TtRecO and TtSSB to the 5’ biotinylated ssDNA were measured by biolayer interferometry using an Octet RED96 instrument (ForteBio). The buffer used for the association and dissociation comprised of 20 mM HEPES pH 7.5, 150 mM NaCl, 0.01 % Tween 20 and 0.1 % BSA. 2 M MgCl_2_ was used for final regeneration.

## Results and Discussion

### Sequence and structure analysis of TtRecF protein

TtRecF is crystallized in orthorhombic space group (P22_1_2_1_) with one molecule in the asymmetric unit (Table 1). RecF exists as a monomer in solution (Koroleva *et al*., 2007). The RecF protein has two domains *viz* ATPase domain and DNA binding domain (Fig. 1a). The RecF protein share a strong homology with Rad50 and structural maintenance of chromosome (SMC) proteins found in eukaryotes and archaea. Both the proteins are involved in DNA repair activities (Rojowska *et al*., 2014; Lehmann, 2005). The ATPase domain shares a significant structural similarity with Lobe I subdomain of the Rad50 head domain and SMC proteins. ATPase domain consists of α-helix sandwiched between parallel β-sheets from the top and anti-parallel β-sheets from the bottom (Fig. 1a). A similar pattern is present in the ATPase domain of both Rad50 and SMC proteins. The second, DNA binding domain of RecF is mostly α-helical and shares similarity with the Lobe II domain of Rad50 head domain (Supplementary Fig. S2). The RecF protein consists of Walker A, Walker B, D-loop and ABC signature motifs (Fig. 1a). The Walker A motif is an ATP binding loop with the consensus sequence, GXXGXGKST. It interacts with α- and β-phosphates of di- and tri-nucleosides. The ABC signature motif is involved in sensing the γ-phosphate of tri-nucleosides (Moncalian *et al*., 2004). The Walker B motif consists of four aliphatic amino acids followed by two negatively charged residues. The negatively charged residue coordinates with the Mg^2+^ ion or polarizes the attacking water molecule (Hanson & Whiteheart, 2005) and the D-loop consists of an invariant aspartic acid of unknown function. Structure based sequence alignment of RecF proteins shows that these motifs are conserved across the organisms (Fig. 2).

**Figure 1.**
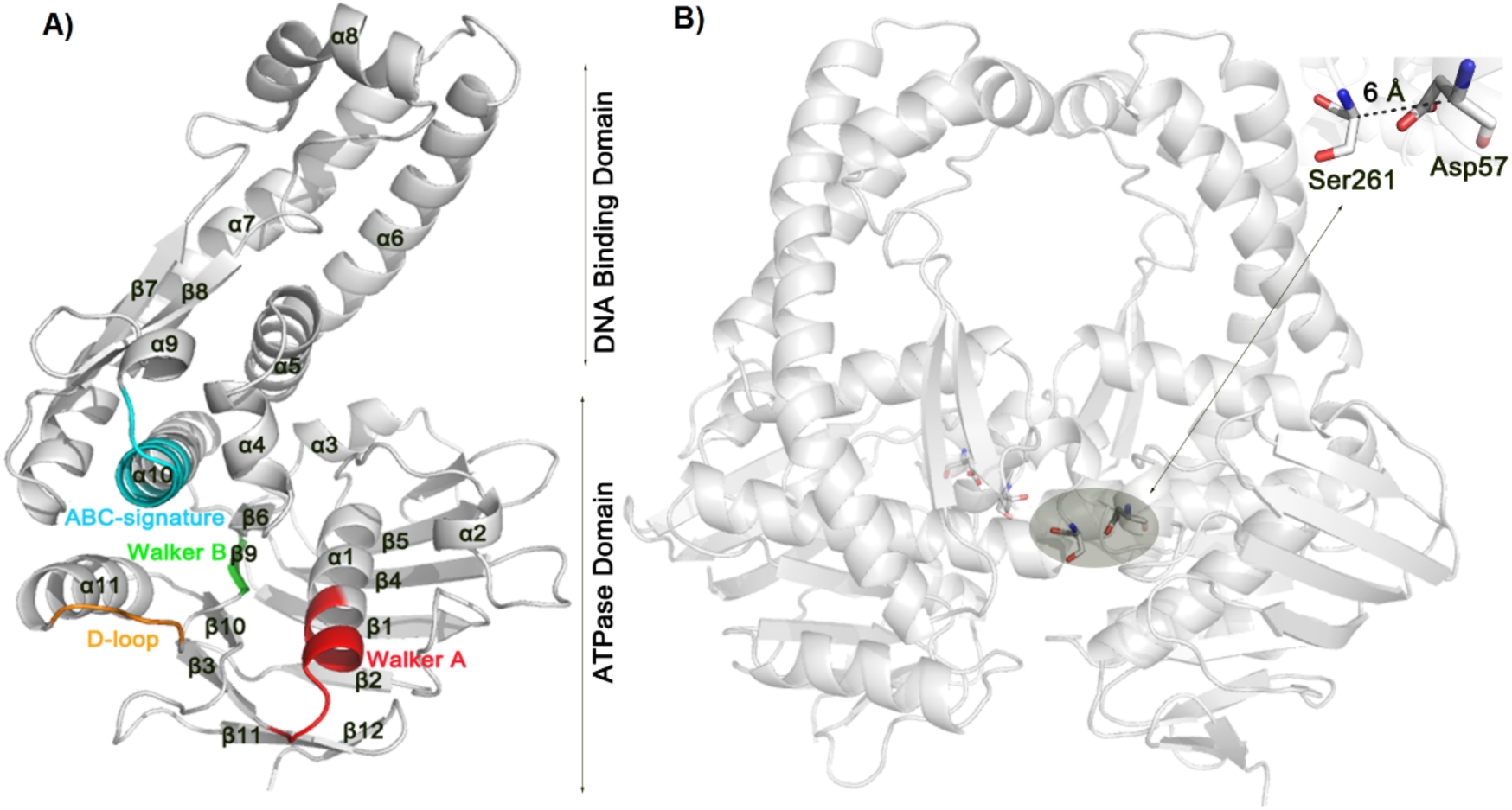
Crystal structure of RecF in monomeric and its modelled dimeric state. **(A)** Cartoon representation of RecF with individual motifs labelled in distinct colors. The ATPase and DNA binding domains are represented at the side of the structure. **(B)** The dimeric interface of the RecF was modelled using Rosetta server. The Cα atom of the residues Ser261 and Asp57 are present at a distance of 6 Å. These residues were selected for mutation to cysteine residues (TtRecF_D57C_ and TtRecF_S261C_).

**Figure 2.**
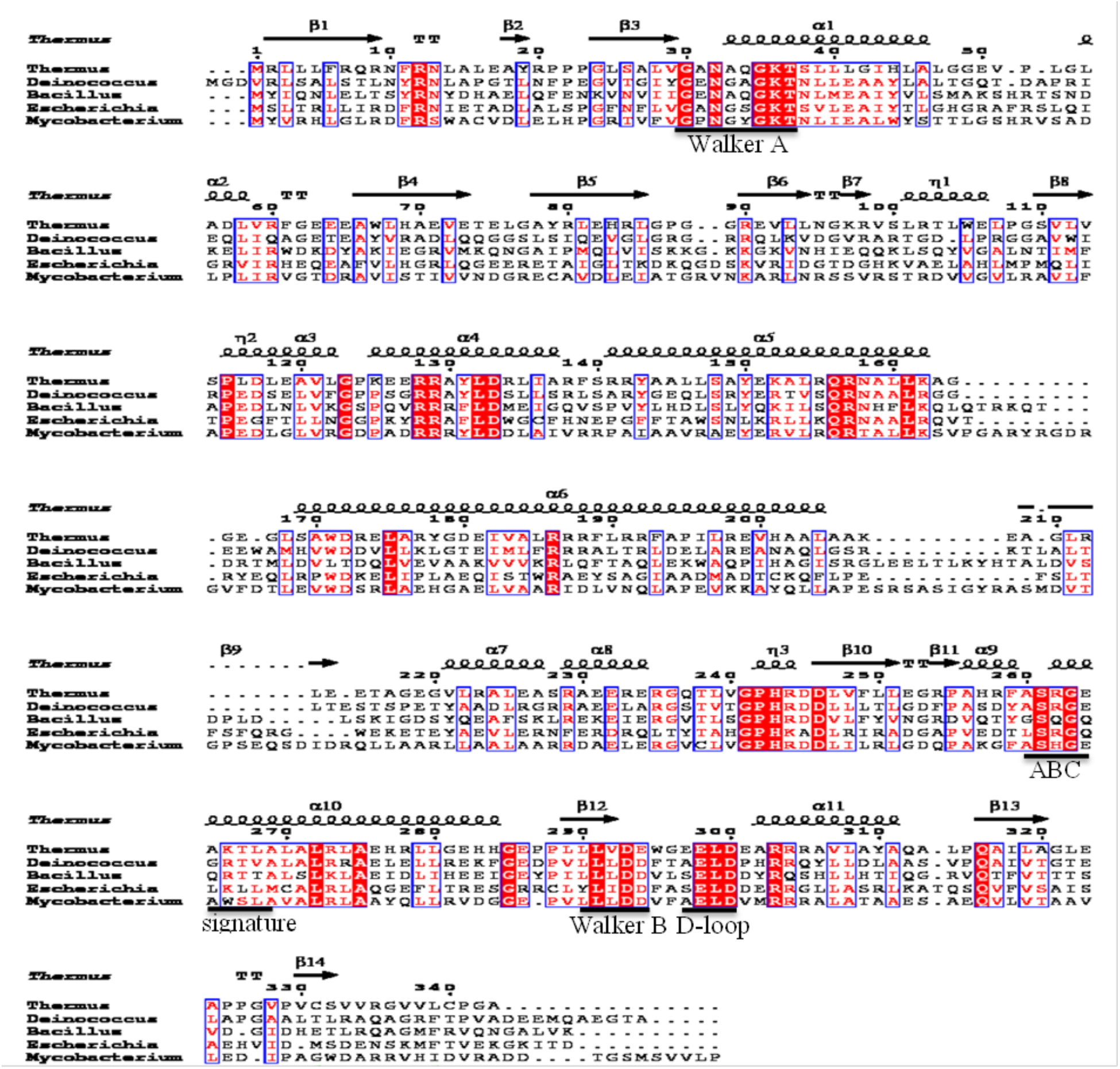
Structure based multiple sequence alignment of RecF proteins. The RecF sequences are taken from various organisms (marked at the beginning). Highly conserved regions are marked in red and their names are mentioned at the base of the sequence.

**Table 1.** Data processing and refinement statistics of TtRecF and TtRecR.

| **Protein name** | **TtRecF** | **TtRecR** |
| --- | --- | --- |
| **Wavelength (Å)** | 0.95 | 1.54 |
| **Resolution limit (Å)** | 51.12-1.58<br>(1.60-1.58) | 50.11-2.50<br>(2.60-2.50) |
| **Unit cell dimensions (Å)** | 37.70<br>74.38<br>153.35 | 35.92<br>100.22<br>136.02 |
| **R<sub>merge</sub> (%)** | 4.4(35.5) | 6.7(89.5) |
| **# of Unique reflections** | 60237 | 8602 |
| **# of molecules in the asymmetric unit** | 1 | 1 |
| **Completeness (%)** | 99.3(82.0) | 96.8(93.6) |
| **Multiplicity** | 4.8(3.8) | 10.3(9.1) |
| **Mean((I)/sd(I))** | 17.2(3.5) | 14.6(1.5) |
| **Refinement statistics** |  |  |
| **R<sub>cryst</sub>/R<sub>free</sub> (%)** | 17.0/22.0 | 23.1/28.0 |
| **rmsd bond length (Å)** | 0.019 | 0.007 |
| **rmsd bond angle (°)** | 1.97 | 1.20 |
| **Average B-factors (Å<sup>2</sup>)** |  |  |
| **Protein** | 31.58 | 76.78 |
| **Water** | 38.17 | 63.65 |
| **Ligands** | - | 65.48 |
| **Ramachandran statistics (%)** |  |  |
| **Favored region** | 97.3 | 97.4 |
| **Allowed region** | 2.4 | 2.1 |
| **Outliers** | 0.3 | 0.5 |
| **PDB ID** | **5ZWU** | **5ZVQ** |

*The data collection statistics values for the highest resolution shell are given in parentheses*

### Dimerization of TtRecF protein

RecF shows ATP dependent dimerization and dimer dependent DNA binding (Koroleva *et al*., 2007). Recently, the dimeric structure of RecF from *Thermoanaerobacter tengcongensis* (Tte) has been reported in complex with ATP and ATPγS (PDB ids: 5Z68 and 5Z69). The dimeric interface of TteRecF in both the complexes are very different and the dimeric state in complex with ATP is regarded as the biologically relevant complex (Tang *et al*., 2018). The ATP bound dimeric TteRecF was used to model the dimeric interface of TtRecF. The dimeric TtRecF model was further refined and energy minimized using Rosetta docking refinement server (Fig. 1b) (Lyskov & Gray, 2008; Moretti *et al*., 2018). The dimeric interface area of TtRecF was higher than that of the TteRecF. No conserved hydrogen bonds were found in any of the two dimers. However, they commonly share regions such as, ABC signature and Walker A motifs present at their dimeric interface. The percentage contribution of polar and non-polar interactions for the stabilization of dimeric interface in both the proteins are similar (Supplementary Fig. S3a and Supplementary Table S2).

The dimeric state of RecF is the prerequisite for binding to the dsDNA or at the junction of dsDNA-ssDNA (Morimatsu & Kowalczykowski, 2003). The RecF shows minimal ATPase activity and its dimeric state might get stabilized at high concentrations (2mM) of ATP (Michel-Marks *et al*., 2010; Koroleva *et al*., 2007). The slow ATPase activity may support RecF for longer binding time at the dsDNA and may help in scanning the dsDNA for the break site. Unsurprisingly, its role in DNA repair still remains elusive. One way to characterize its roles is by stabilizing its dimeric interface. Thus, the modeled dimeric TtRecF interface residues, Asp57 and Ser261 were selected for mutation to cysteine residues (TtRecF_D57C_ and TtRecF_S261C_) (Fig 1b). The TtRecF mutants (ttRecF_D57C_ and ttRecF_S261C_) were purified in soluble fraction. The wild type TtRecF exits as a monomer in solution and formed dimer in the presence of ATP (Koroleva *et al*., 2007; Tang *et al*., 2018). Thus, it was expected that the cysteine residues would come closer in the presence of ATP molecules and form a disulfide bond in an oxidative environment resulting in a stable dimeric TtRecF. To our surprise, the purification of both the TtRecF cysteine mutants (ttRecF_D57C_ and ttRecF_S261C_) resulted in a non-canonical dimeric, as well as, monomeric population in solution. The dimeric populations of the mutant proteins are reduced to monomeric state after treatment with a reducing agent (Fig. 3). As mentioned earlier, the TteRecF shows that the two monomers can exist in totally different orientations in the presence of non-hydrolysable ATP (Aravind *et al*., 1998). This suggests that the dimeric orientations of RecF might depend on its surrounding environment (Supplementary Fig. 3b). Most importantly, ABC ATPases and DrRecF showed ATP dependent dimerization at high ATP and protein concentrations (Leiros *et al*., 2005; Koroleva *et al*., 2007). In contrast, TtRecF_D57C_ and TtRecF_S261C_ mutants formed di-sulphide bridges in the absence of ATP. Several attempts to crystallize the mutants remain unsuccessful. Coupled enzyme and calorimetric assays (using malachite green) were performed on the wild type TtRecF and its mutants to monitor the changes in the ATPase activity. These experiments showed negligible ATPase activity for the wild type as well as mutant TtRecF proteins, both in the presence and absence of dsDNA/gap DNA (gDNA). A more sensitive ATPase assay (radioactivity assay) might be helpful in determining the changes in ATPase activity.

**Figure 3.**
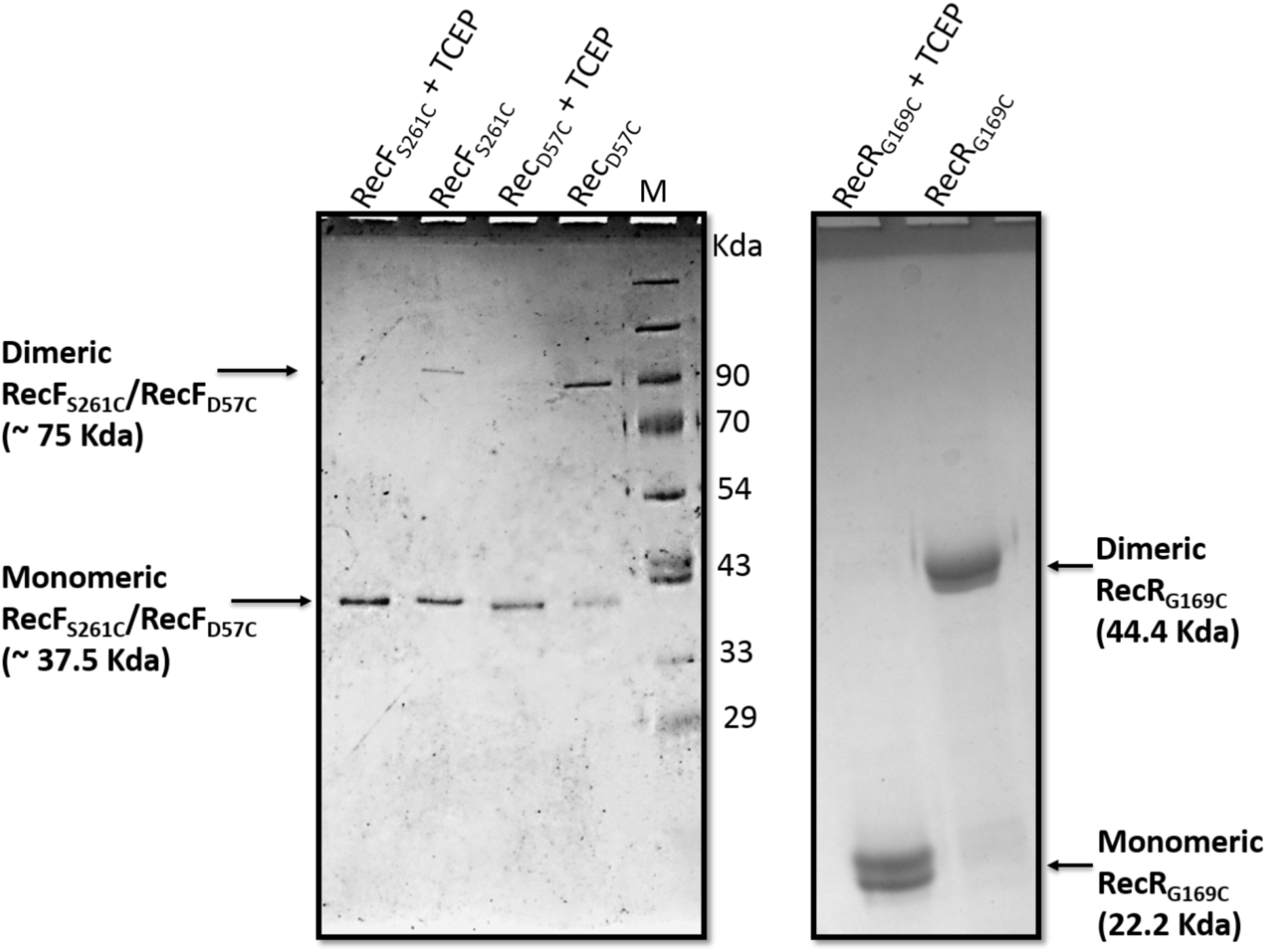
SDS-PAGE analysis of TtRecF_D57C_, TtRecF_S261C_ and TtRecR_G169C_ proteins upon treatment with TCEP (tris(2-carboxyethyl)phosphine). TtRecF_D57C_, TtRecF_S261C_

### Structural analysis of TtRecR protein

RecR from *Thermus thermophilus* (TtRecR) was purified and crystallized in I222 space group. The RecR structure from *D. radiodurans* (DrRecR) (PDB -id: 1VDD) was used as a search model for Molecular Replacement (MR) calculations (Table 1). The model was successfully generated for TtRecR in I222 space group. The TtRecR protein solved in I222 space group consists of one molecule in the asymmetric unit (Fig. 4a). Interestingly, a tetrameric assembly of the RecR protein was generated by combining the other three crystal symmetry mates. The tetrameric assembly of monomeric subunits ensued to a ring shaped structure with a central cavity. The dimensions of the central cavity are 30 and 35 Å, large enough to pass a dsDNA (Fig. 4b).

**Figure 4.**
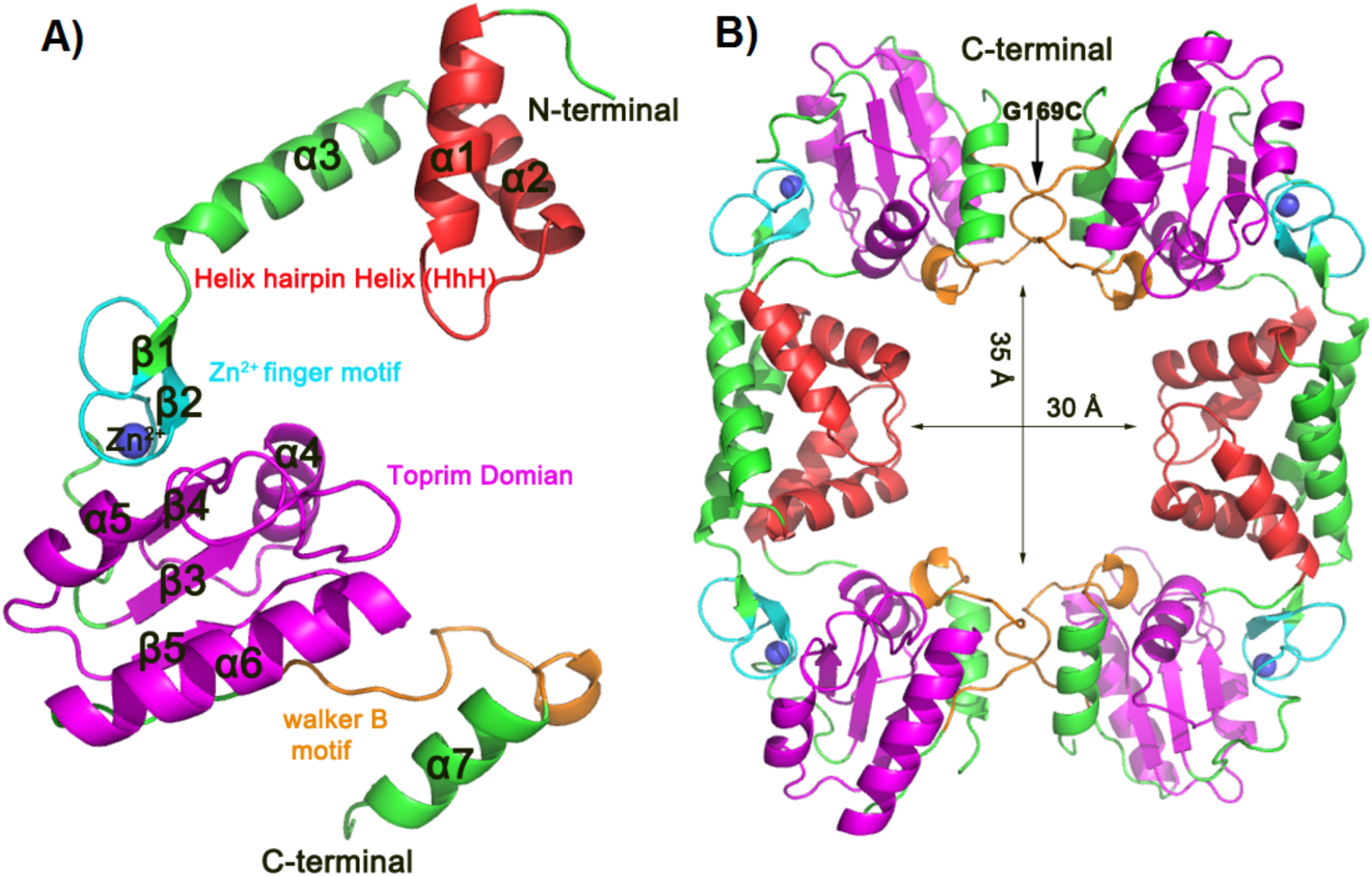
Monomeric and tetrameric assemblies of the wild-type TtRecR protein. **(A**) Crystal structure of the wild-type ttRecR. The motifs in the structures are marked in different colors and their names are represented in similar colors. The blue colored sphere represents a Zinc ion. **(B)** A tetrameric assembly of the wild-type ttRecR is generated by combining the symmetry mates. The N- and C-terminals of the protein are involved in a domain swap and form a stable tetrameric assembly. The tetrameric assembly creates a central hole of 30-35 Å. The Gly169 present at the C-terminal of TtRecR protein was mutated to cysteine residue (TtRecR_G169C_).

The RecR protein consists of a Helix hairpin Helix (HhH) motif, a Cys4 zinc-finger motif, a Toprim domain and a Walker-B motif (Fig. 4a). These motifs are highly conserved across prokaryotes (Fig. 5). Helix hairpin Helix (HhH) motif is present at the N-terminal of the protein. It has a consensus sequence, hxxhxGhGxxxAxxhh (where, h is any hydrophobic residue and x could be any amino acid). The motif comprises two α-helices joined by a hairpin loop. HhH motifs are commonly found in many of the DNA replication and repair proteins, such as Endonuclease III of *E. coli*, human DNA polymerase β and Adenine DNA glycosylase (Doherty *et al*., 1996; Thayer *et al*., 1995). In RecR proteins, the HhH motifs are domain swapped with the other adjacent molecule in the crystal to form a dimeric structure. The conserved Lys23, Arg27 and the loop containing Leu17, Pro18, Gly19 and Ile20 are projected towards the central cavity of the tetrameric TtRecR structure (Honda *et al*., 2008; Honda *et al*., 2006). These residues might play an important role in DNA binding. The mutation of Lys23 and Arg27 at equivalent positions in DrRecR resulted in reduced DNA binding (Lee *et al*., 2004).

**Figure 5.**
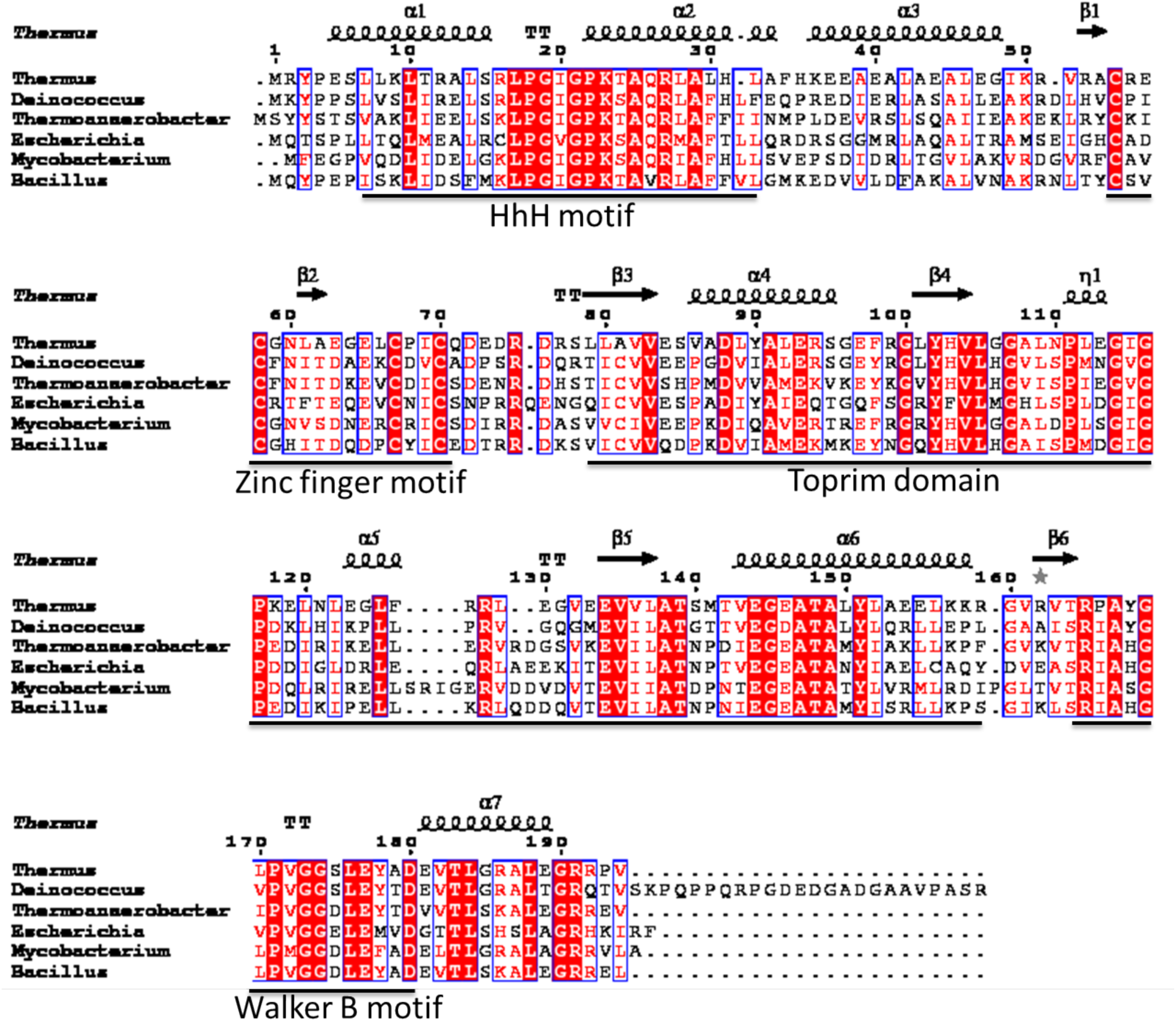
Structure based sequence alignment of RecR proteins. The RecR sequences are taken from various organisms (marked at the beginning of sequences). Highly conserved regions are marked in red and motifs are underlined with their names marked the base.

TtRecR comprised of four conserved cysteine residues, which coordinate with the Zn^2+^ ion to form a Cys4 zinc finger motif (Fig. 4a). The zinc finger motif in proteins is found to be important for the protein-protein and protein-DNA interactions (Kusakabe *et al*., 1999). The zinc finger is formed by a combination of two antiparallel β-strands and a α-helix. In DNA binding proteins, the α-helix binds to the major groove of the DNA. However, the Cys4 zinc-finger motif in RecR plays an important role in maintaining its three-dimensional structure. The mutation of individual cysteine residues has an effect on the protein stability; however, its interaction with the DNA or RecO protein remains unchanged (Tang *et al*., 2014).

The toprim domain consists of four stranded parallel β-sheet flanked by a α-helix from one side and two α-helices from the other side (Fig. 4a). This represents a typical Rossmann type nucleotide-binding fold. The toprim domain is also present in DNA binding proteins such as Topoisomerase (types IA and II), DnaG-type primases and old family nucleases. The toprim domain comprised of a conserved glutamic acid residue at the start of the toprim domain and a DxD motif towards the end of the domain. The glutamic acid may act as a general base in nucleotide polymerization by primase and stand ligation by topoisomerase, whereas in case of strand cleavage by topoisomerase and nuclease, as a general acid (Aravind *et al*., 1998). The DxD motif is required for the binding of the Mg^2+^ ion and is important for the function of all toprim-containing enzymes. The glutamic acid was found to be partially conserved and DxD motif is absent in RecR proteins (Allemand *et al*., 2005; Aravind *et al*., 1998; Fu *et al*., 2009). Thus, the toprim domain of RecR may not act as those seen in topoisomerase and primase enzymes. However, it can perform an important role in DNA binding and recombination mediator protein interactions.

The Walker-B motif and the preceding α-helix take part in domain swap at the C-terminal and help in forming a tetramer (Fig. 4b). The C-terminal deleted mutants of DrRecR showed aggregation during purification (Lee *et al*., 2004), whereas TtRecR showed a dimeric population. However, the N-terminal deletion of TtRecR was a monomer. This showed that it is the N-terminal and not the C-terminal that is required for stable dimer formation (Honda *et al*., 2008; Tang *et al*., 2012).

### Tetramerization of TtRecR protein

Both TtRecR and EcRecR are known to form dimers in solution. However, a homo-tetrameric assembly was observed in the crystal structures of TtRecR, DrRecR and TteRecR. The RecR protein interacts with RecF and RecO proteins in tetrameric state (Honda *et al*., 2008). However, the factors responsible for the tetramerization of RecR are not known. Also, the interactions of RecR with other cognate proteins for binding to the DNA are not clear. Earlier studies showed that the N-terminal of RecR is required for stable dimer formation and not the C-terminal (Honda *et al*., 2006). Thus, the swap region of the C-terminal was targeted for cysteine mutation to form a stable tetramer in solution. Gly169 present at the C-terminus of TtRecR is mutated to cysteine (Fig. 4b). The purification of TtRecR_G169C_ mutant resulted in a dimeric 44.4 kDa band along with a small quantity of monomeric band at 22.2 kDa (Fig. 3). This concludes that the TtRecR_G169C_ mutant exists as a tetramer in solution even at very low concentration. The stable tetrameric TtRecR has been tested for its interaction with ssDNA and other repair mediator proteins.

### Stabilized tetrameric mutant of TtRecR shows distinct interaction properties

The RecR protein interacts with both the RecF and RecO proteins by forming either RecOR or RecFR complexes. The structural studies on RecOR complex from *D. radiodurans* showed that the complex consists of a heterohexamer in a ratio of 4RecR:2RecO (PDB id - 4JCV) (Timmins *et al*., 2007). The tetrameric DrRecR is bound with two monomeric RecO proteins at the adjacent sides of its central cavity. The OB-fold domain of the RecO protein is pointed towards the interior of the DrRecR cavity. The DrRecOR complex shows preferential binding towards the 3’ overhang of ssDNA-dsDNA hybrid. The site directed mutagenesis studies on DrRecR and DrRecO showed that the conserved Glu146 residue present in the toprim domain of DrRecR (equivalent residue; Glu144 in TtRecR) and His93 of DrRecO (equivalent residue not present in TtRecO) are essential for the formation of this complex (Timmins *et al*., 2007). Structural and biochemical studies on the RecOR complex from *T. tengcongensis* also showed that the loop region (106 to 121) present in the toprim domain of RecR is essential for RecO binding (Tang *et al*., 2012).

RecR can also form a complex with the RecF protein. The TtRecFR complexes are formed in 4:2 ratio (4TtRecR:2ttRecF) (Honda *et al*., 2008). Both, the N-terminal and the C-terminal of TtRecR are required for the TtRecFR complex formation. The TtRecR interacts with the TtRecF in a tetrameric state. The SASX data suggests that the assembly of this hetero-hexameric complex is such that the two RecF can interact with tetrameric TtRecR molecules, either on the opposite sides of the central RecR cavity or from the same side. However, in both the cases, the central cavity of the tetrameric TtRecR was covered by the TtRecF proteins, leaving no space for DNA to pass (Honda *et al*., 2008). The mutation of Glu144 residue in TtRecR showed reduced binding to TtRecO protein (as also observed in the equivalent mutation in DrRecR) and completely abolished its interaction with TtRecF. Thus, both (TtRecF and TtRecO) proteins share an overlapping binding site for their binding to the TtRecR protein and both the proteins interacts with the tetrameric RecR (Honda *et al*., 2006).

To elucidate the effect of the TtRecR tetramerization (TtRecR_G169C_) on the binding of the wild type TtRecF and TtRecO proteins, Native-PAGE and EMSA experiments were performed. The proteins TtRecF and TtRecO have higher pI values of 8.92 and 10.25, respectively as compared to RecR (pI 5.43). Thus, TtRecF and TtRecO were not able to move in the native-PAGE. An additional band appeared, when TtRecR and TtRecR_G169C_ were incubated with the TtRecF protein, suggesting a stable TtRecFR/TtRecFR_G169C_ complex formation in solution (Figs. 6a and 6b). Nevertheless, the TtRecOR complex formation should not be neglected. The TtRecR/TtRecR_G169C_interactions may not be able to subside the positive charge on the TtRecO protein and hence the complex remains in the well during the electrophoresis. This suggest that the interactions of TtRecF and TttRecO proteins with the wild type and mutant TtRecR would be similar. Further, these interactions were also studied in the presence of fluorescein (FAM) labelled ssDNA. The incubation of TtRecF, TtRecO, TtRecR and mutant TtRecR_G169C_ proteins do not result in any band shift. A shift in the ssDNA band was observed when the TtRecO or TtRecF proteins were incubated with the wild type TtRecR, whereas, these shifts were missing in the presence of the mutant, TtRecR_G169C_ (Figs. 7a and 7b).

**Figure 6.**
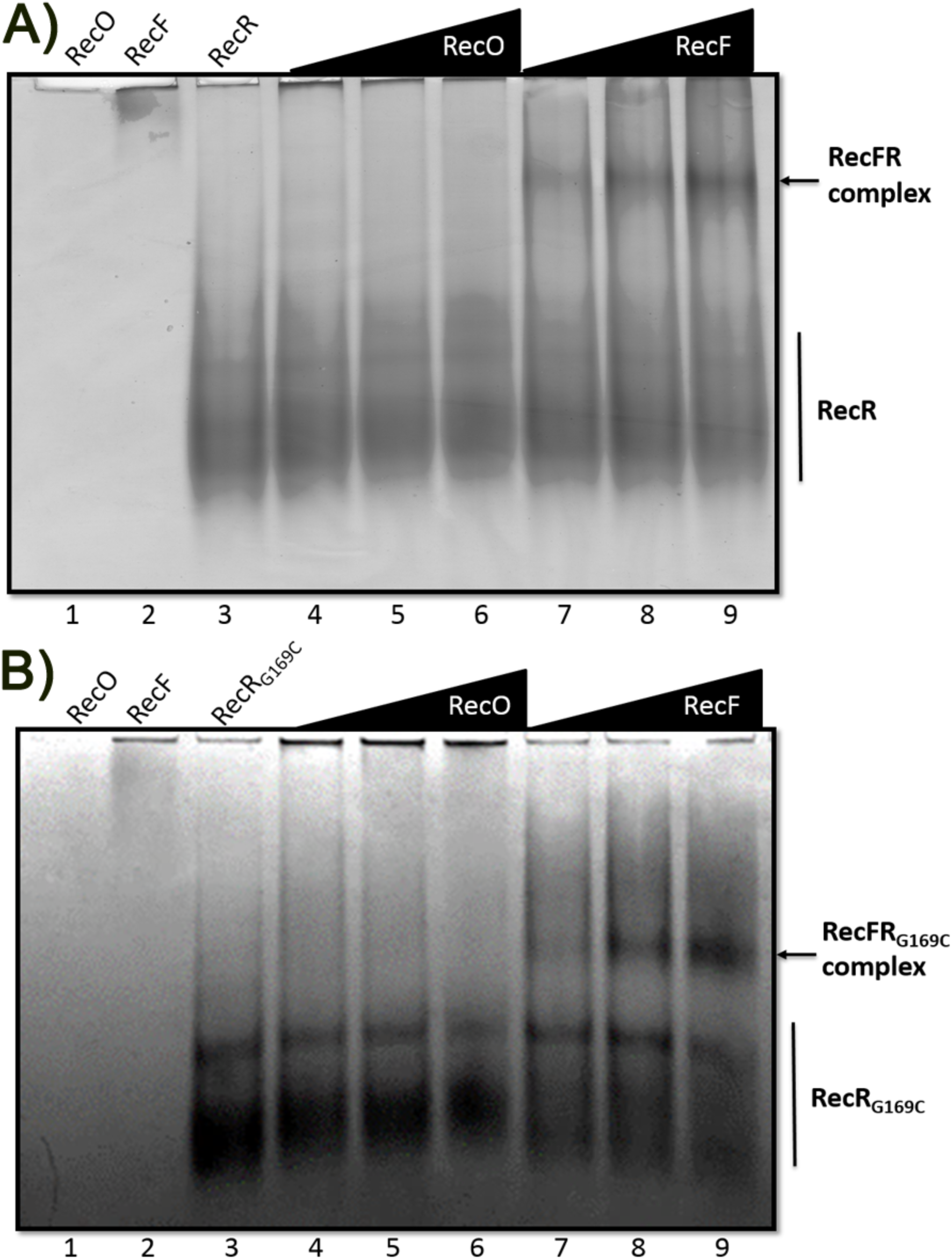
Analysis of protein-protein interactions RecF and RecO with the wild type RecR and mutant ttRecR_G169C_ using native-PAGE. The higher pI values of RecO (pI – 10.3) and RecF (pI – 8.3) restrict its mobility in the native-PAGE (lanes 1-2), whereas the mobility of RecR/RecR_G169C_ is facilitated by lower pI value (pI – 6.3) (lane 3). 10 µM of RecO, RecF and RecR/RecR_G169C_ was loaded in the lanes 1-3. **(A)** 10 µM of RecR was incubated with 10, 20 and 40 µM concentrations of RecO (lanes 4-6) and with 10, 20 and 40 µM concentrations of RecF (lanes 7-9). **(B)** 10 µM of mutant RecR_G169C_ was incubated with 10, 20 and 40 µM concentrations of RecO (lanes 4-6) and with 10, 20 and 40 µM concentrations of RecF (lanes 7-9).

**Figure 7.**
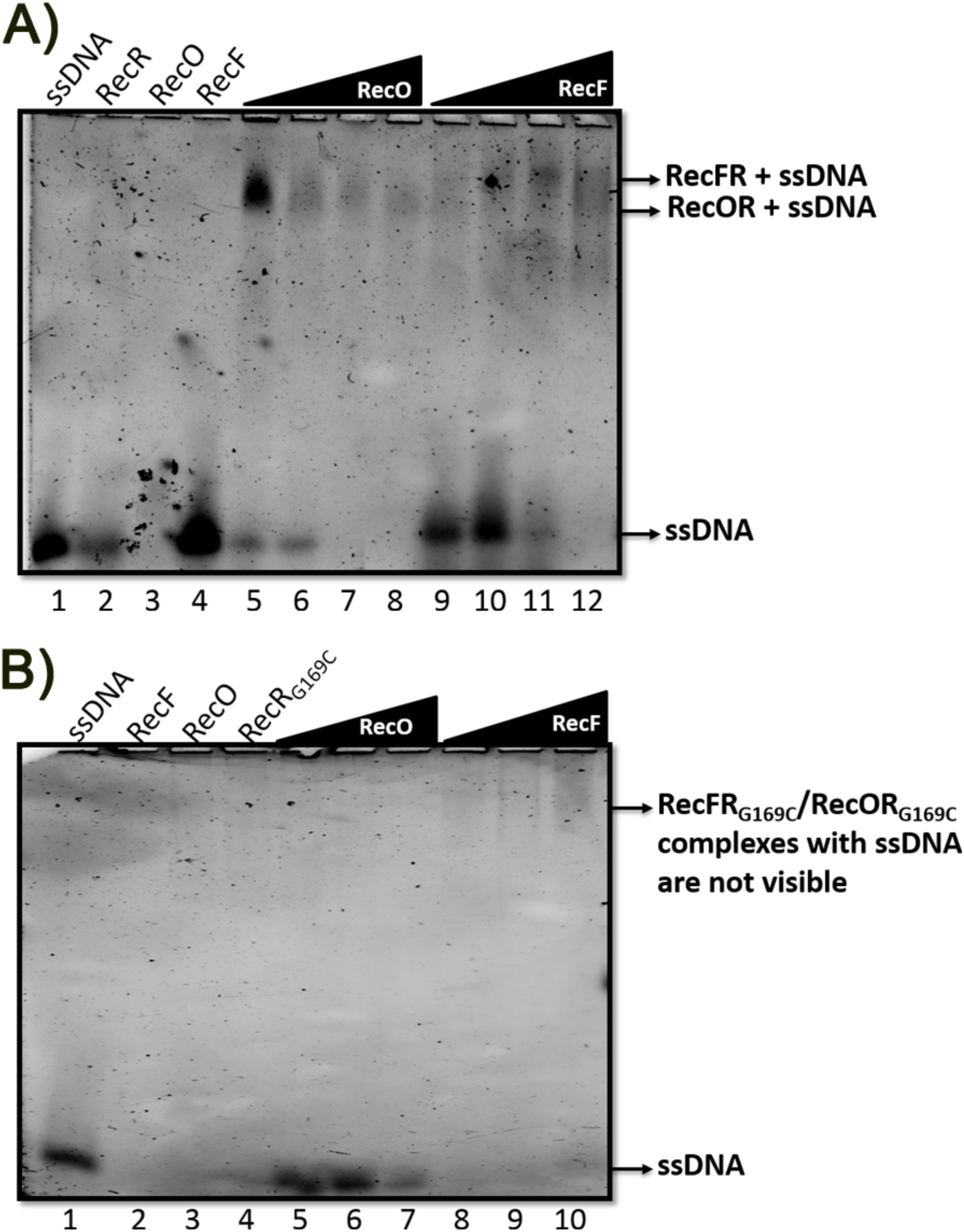
Analysis of protein-protein and protein-ssDNA interactions of RecF and RecO with the wild type RecR and mutant ttRecR_G169C_ using EMSA. Fluorescein (FAM) labelled dT_35_ was used as ssDNA for the interaction studies. **(A)** ssDNA was incubated with the 10 µM of RecR, RecO and RecF (lanes 2-4). ssDNA was incubated with 10 µM of RecR and 5, 10, 20 and 40 µM of RecO (lanes 5-8). ssDNA was incubated with 10 µM of RecR and 5, 10, 20 and 40 µM of RecF (lanes 9-12). **(B)** ssDNA was incubated with the 10 µM of RecF, RecO and RecR_G169C_ (lanes 2-4). ssDNA was incubated with 10 µM of RecR_G169C_ and 5, 10 and 20 µM of RecO (lanes 5-7). ssDNA was incubated with 10 µM of RecR_G169C_ and 5, 10 and 20 µM of RecF (lanes 8-10).

### TtRecO displays higher binding to ssDNA in the presence of the wild type TtRecR

The individual binding affinities of the wild type and mutant proteins with ssDNA have been monitored using Bio-Layer Interferometry (BLI). The wild type TtRecR exhibited negligible binding with the ssDNA, whereas the mutant TtRecR_G169C_ showed binding and the binding data can be fitted using mass transport binding model (Figs. 8a and 8b). However, this has not helped in determining the K_D_ value accurately. TtRecO showed a strong binding with the ssDNA (Fig. 8c). The binding curve can be fitted in the heterogeneous ligand model, which suggests that the TtRecO binds to ssDNA with two different off-rate constants; hence RecO might have two different binding modes. A tenfold difference has been recorded between the two binding affinities. The N-terminal OB fold domain of the RecO proteins are mainly responsible for DNA binding, however, the C-terminal domain on the protein was also found to have relatively lesser affinity towards the DNA, as observed in DrRecO (Leiros *et al*., 2005). This would be the reason for the biphasic behavior of TtRecO binding.

**Figure 8.**
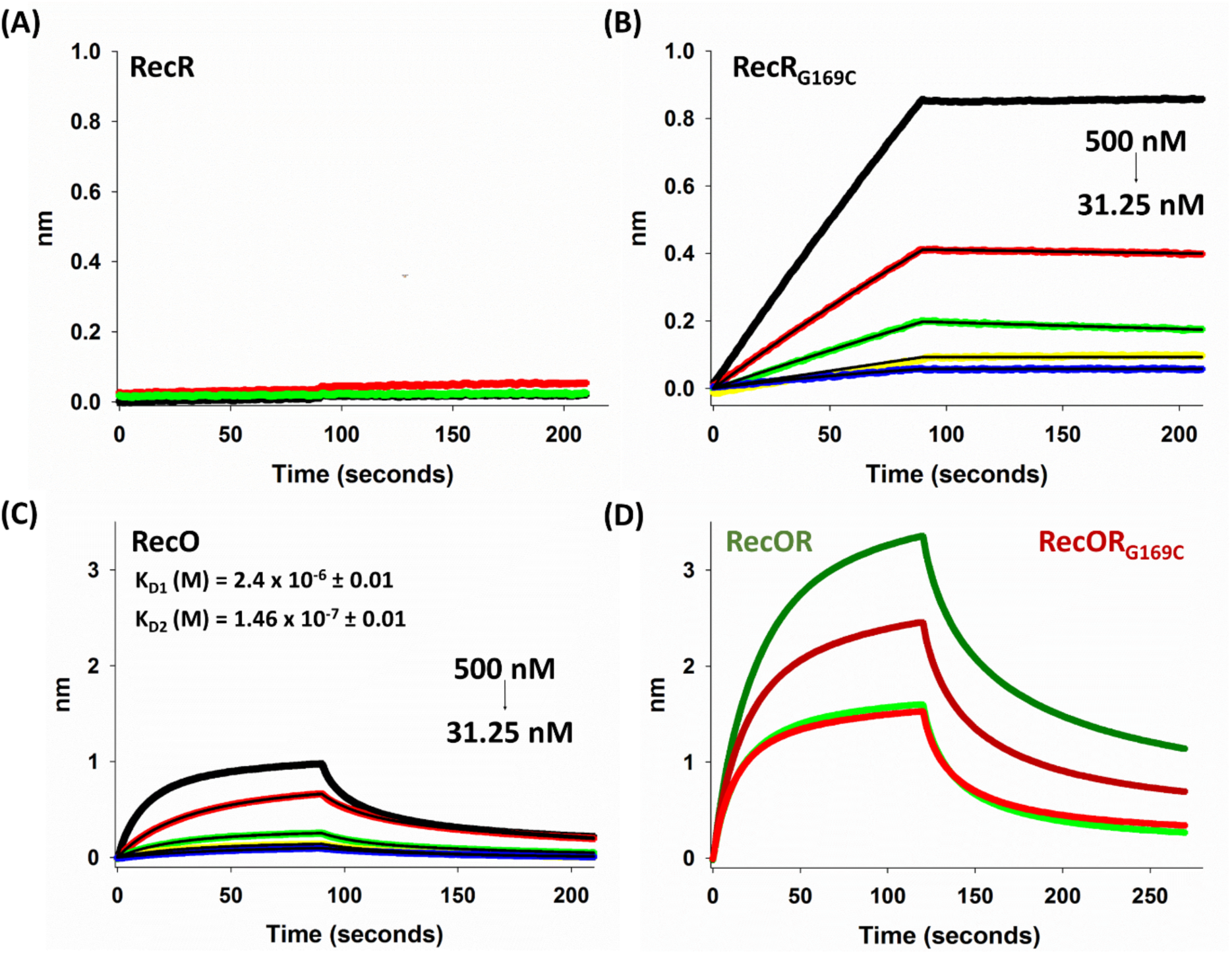
Interactions of TtRecR, TtRecR_G169C_, TtRecO and RecOR/RecOR_G169C_ complexes with ssDNA. **(A)** Interactions of ssDNA with different concentrations of the wild type TtRecR. **(B)** Interactions of ssDNA with different concentrations (colored lines) of the stabilized tetrameric TtRecR_G169C_. The curves are fitted using the mass transport model (black lines). **(C)** Interactions of ssDNA with different concentrations (colored lines) of TtRecO. The curves are fitted using heterogeneous ligand model (Black lines) and the two binding affinities (K_D_) are marked on it. **(D)** Interactions of the ssDNA with the TtRecO in the presence of the wild type TtRecR and mutant TtRecR_G169C_. The dark green curve denotes incubation of ssDNA with 500 nM of TtRecR and 250 nM TtRecO. The light green curve denotes incubation of ssDNA with 250 nM of TtRecR and 250 nM TtRecO. The dark red curve denotes incubation of ssDNA with 500 nM of TtRecR_G169C_ and 250 nM TtRecO. The light red curve denotes incubation of ssDNA with 250 nM of TtRecR_G169C_ and 250 nM TtRecO.

The TtRecO interacts with the TtRecR proteins to form TtRecOR complex in a ratio of 2RecO:4RecR (Timmins *et al*., 2007). Its interaction with the ssDNA was monitored. A huge jump in the nm was observed in the presence of TtRecR and TtRecO (Fig. 8d). This normally means a better binding affinity, however, considering the measurement principle of the instrument, the shift in the interference pattern is the result of changing thickness at the tip of the biosensors. Here, the increase in the thickness would be due to the binding of TtRecOR complex instead of only TtRecO. A higher shift was observed, when the concentration of the wild type TtRecR or the mutant TtRecR_G169C_, was twice the TtRecO concentration, this effect was more pronounced in the case of the wild type TtRecR (Fig. 8d). Indicating, that the conformational flexibility between the dimeric TtRecR protein would assist the binding of TtRecO protein. Moreover, a decrease in the shift is observed, when equal concentrations of the TtRecO and TtRecR/RecR_G169C_ proteins were used and this shift was equivalent to the TtRecO shift (Figs. 8c and 8d). This shows a molarity dependent interaction stabilization between the TtRecR and TtRecO proteins.

### TtRecF mutants shows affinity for ssDNA

The native-PAGE has confirmed the formation of TtRecFR and TtRecFR_G169C_ complexes in the solution. EMSA results showed that TtRecF has no affinity for ssDNA, however a band shift appeared, when incubated with the wild type TtRecR protein. The BLI experiments also complimented these results. Conversely, a slight increase in the nm shift was observed in the presence of the wild type TtRecR and mutant TtRecR_G169C_ (Figs. 9a and 9b). These shifts are perhaps due to mass transfer limited kinetics, where the rate of binding to the target site is faster than its diffusion. This shows that the TtRecFR complex has negligible affinity towards the ssDNA as compared to the TtRecOR complex. The RecFR complex has been shown to bind at the junction of gDNA and needs a free 5’end for proper binding (Morimatsu & Kowalczykowski, 2003). It suggests that the break site could be recognized by the RecFR complex.

**Figure 9.**
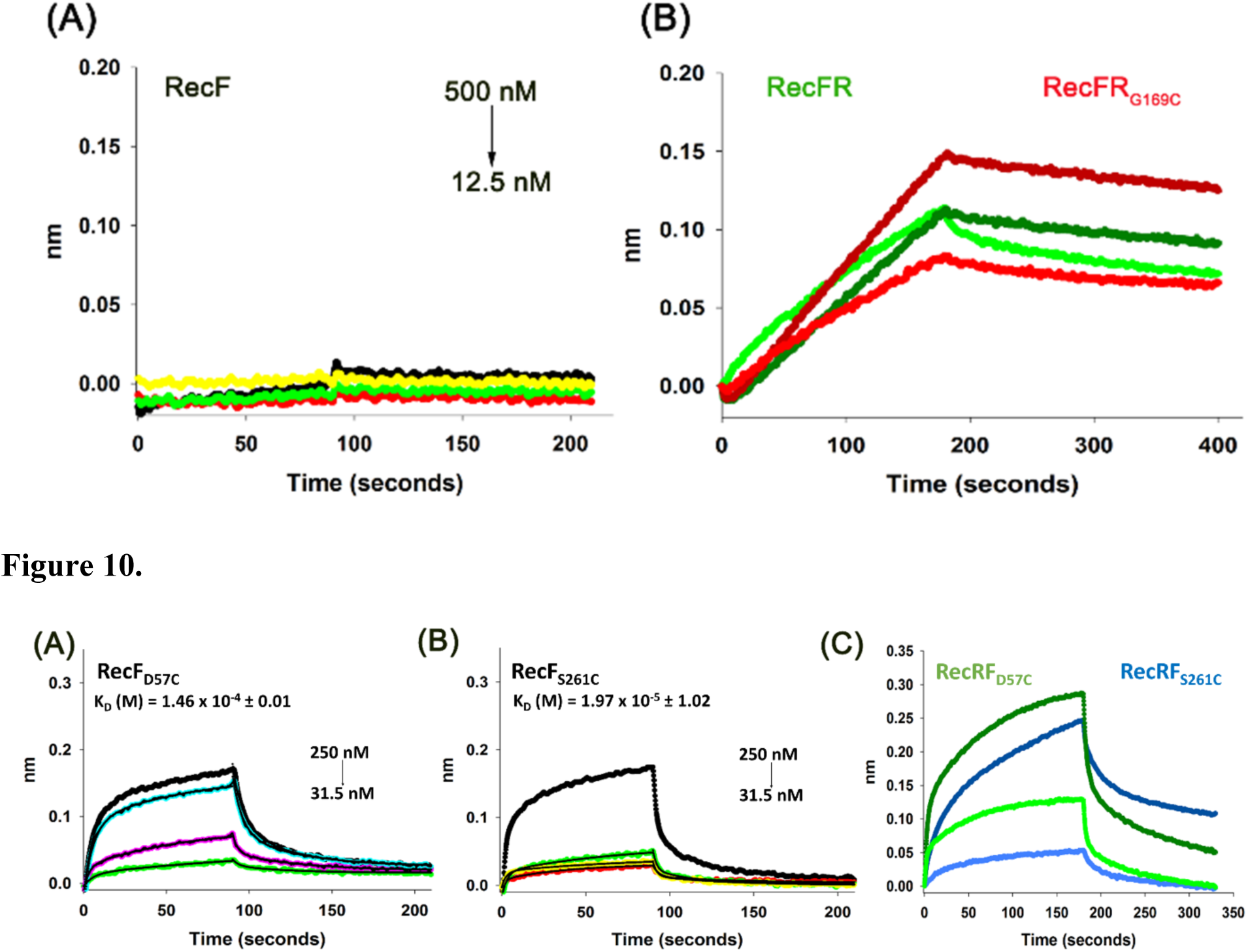
Interactions of TtRecF with ssDNA in the presence and absence of RecR/RecR_G169C_. **(A)** Interactions of ssDNA with different concentrations of the wild type TtRecF. **(B)** Interactions of the ssDNA with the TtRecF in the presence of the wild type TtRecR and mutant TtRecR_G169C_. The dark green curve denotes incubation of ssDNA with 500 nM of TtRecR and 250 nM TtRecF. The light green curve denotes incubation of ssDNA with 250 nM of TtRecR and 250 nM TtRecF. The dark red curve denotes incubation of ssDNA with 500 nM of TtRecR_G169C_ and 250 nM TtRecF. The light red curve denotes incubation of ssDNA with 250 nM of TtRecR_G169C_ and 250 nM TtRecF.

Surprisingly, in contrast to the wild type TtRecF, the non-canonical dimers formed by the cysteine mutations (TtRecF_D57C_ and TtRecF_S261C_), showed binding towards the ssDNA even in the absence of TtRecR (Figs. 10a and 10b). The binding data can be fitted using heterogeneous ligand method; however the two K_D_ values were very similar. Both the mutants (TtRecF_D57C_ and TtRecF_S261C_) showed binding affinities of 146.3 and 19.7 µM, respectively. A shift in the nm was observed after the addition of the wild type TtRecR. The shift was more pronounced when the concentration of TtRecR proteins was twice the concentration of TtRecF_D57C_/TtRecF_S261C_ mutant proteins (Fig. 10c). It complimented the binding ratios of 2RecF:4RecR in the complexes as suggested in an earlier study (Honda *et al*., 2006). In contrast to the wild type TtRecF, these non-canonical dimeric TtRecF_D57C_/TtRecF_S261C_ mutants can bind to only one face of the TtRecR. This suggests that the binding topology of RecF to RecR can regulate the complex affinity towards the ssDNA and dsDNA. The dsDNA binding property of the RecFR complex is important for the DNA break recognition, whereas the physiological significance of the ssDNA binding needs to be explored.

**Figure 10.** Interactions of TtRecF mutants with ssDNA in the presence and absence of the wild type RecR. **(A)** Interactions of ssDNA with different concentrations (colored lines) of TtRecF_D57C_. The curves are fitted using heterogeneous ligand model (black lines) and the binding affinity (K_D_) is marked on it. **(B)** Interactions of ssDNA with different concentrations (colored lines) of TtRecF_S261C_. The curves are fitted using heterogeneous ligand model (black lines) and the binding affinity (K_D_) is marked on it. **(C)** Interactions of the ssDNA with the TtRecF_D57C_ and TtRecF_S261C_ in the presence of the wild type TtRecR. The dark green curve denotes incubation of ssDNA with 500 nM of TtRecR and 250 nM TtRecF_D57C_. The light green curve denotes incubation of ssDNA with 250 nM of TtRecR and 250 nM TtRecF_D57C_. The dark blue curve denotes incubation of ssDNA with 500 nM of TtRecR and 250 nM TtRecF_S261C_. The light blue curve denotes incubation of ssDNA with 250 nM of TtRecR and 250 nM TtRecF_S261C_.

### TtRecO shows higher affinity for TtSSB bound ssDNA in the presence of tetrameric TtRecR

As soon as the break is formed in the DNA, the helicase and endonuclease activities of RecQ and RecJ, respectively, creates a 3’ overhang. This 3’ overhang is immediately coated by the SSB proteins. The binding affinity (K_D_) of the TtSSB proteins for the ssDNA is ∼ 7.2 nM (Fig. 11), which is approximately thousand times higher than that of the TtRecO protein (2.4 μM). Upon performing the TtSSB association step, a minor dissociation was observed. Strong binding property of TtSSB to ssDNA was further used to monitor the binding of other RecFOR pathway proteins on the TtSSB coated ssDNA and another step of association and dissociation was added to the experiments.

**Figure 11.**
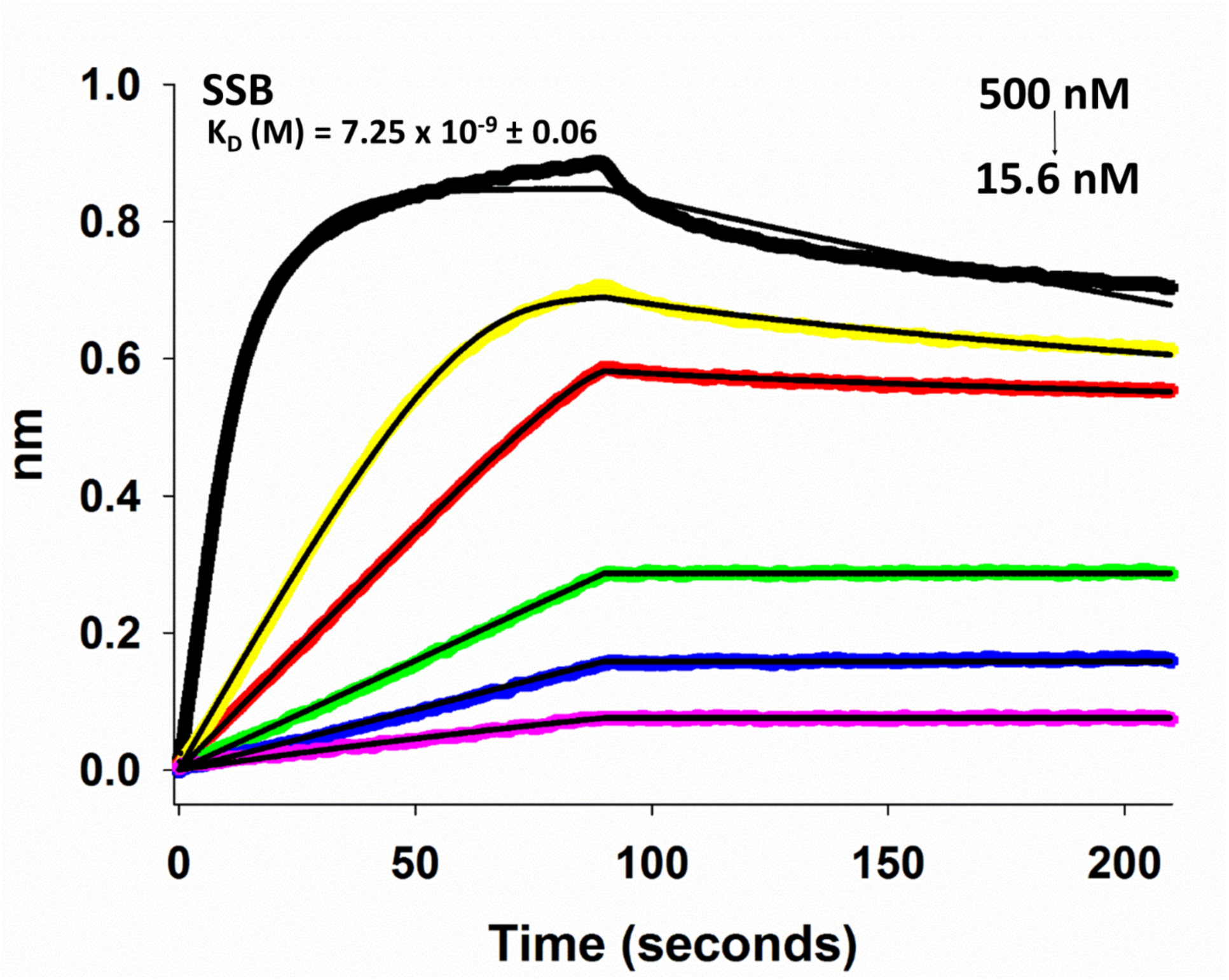
Interactions of single strand binding protein (SSB) with ssDNA. Interactions of ssDNA with different concentrations of SSB are represented by colored curves. The curves are fitted using 1:1 binding model (black lines) and the binding affinity (K_D_) is marked on it.

The wild type TtRecR, TtRecR_G169C_ and TtRecFR complex showed a negligible association to the TtSSB bound ssDNA (Fig. 12a and 12b). However, when the TtSSB coated ssDNA was incubated with the TtRecO protein, an increase in the nm shift was observed (Fig. 12c). This indicates the binding of the TtRecO protein on the TtSSB bound ssDNA. Structural studies on the EcSSB showed that the SSB C-terminal (SSB-Ct) has affinity for the EcRecO protein, which in turn help in its recruitment on the SSB coated ssDNA (Inoue *et al*., 2011). The C-terminal sequence of TtSSB (“EEELPF”) is similar to the EcSSB (“DDDIPF”). Both the TtSSB and TtRecO proteins carry OB folds, required for DNA binding, however, the dimeric TtSSB carry four OB folds as compared to only one OB fold in the monomeric TtRecO. Thus, the binding of two TtRecO proteins at both the C-termini of the TtSSB may result in the decrease in the affinity of the TtSSB protein with the ssDNA. An increased nm shift was observed, when the SSB bound ssDNA was incubated with the TtRecO and TtRecR/TtRecR_G169C_ proteins (Fig. 12d). The nm shift was more pronounced when the concentration of TtRecR/TtRecR_G169C_ proteins was twice than that of the TtRecO concentration. In contrast to wild type TtRecR binding to TtSSB coated ssDNA, the mutant tetrameric TtRecR_G169C_ showed a higher nm shift in the presence of TtRecO. This suggests that the tetrameric state of TtRecR in complex with TtRecO has preferential binding towards the TtSSB coated ssDNA. However, binding of these complexes did not lead to the release of TtSSB from ssDNA. Nevertheless, the TtRecOR complex may reduce the binding affinity of the TtSSB with ssDNA and help in the recruitment of TtRecA protein. The TtRecA proteins forms a nucleoprotein filament on the ssDNA by displacing the SSB and RecOR complex and prepares the DNA for strand exchange.

**Figure 12.**
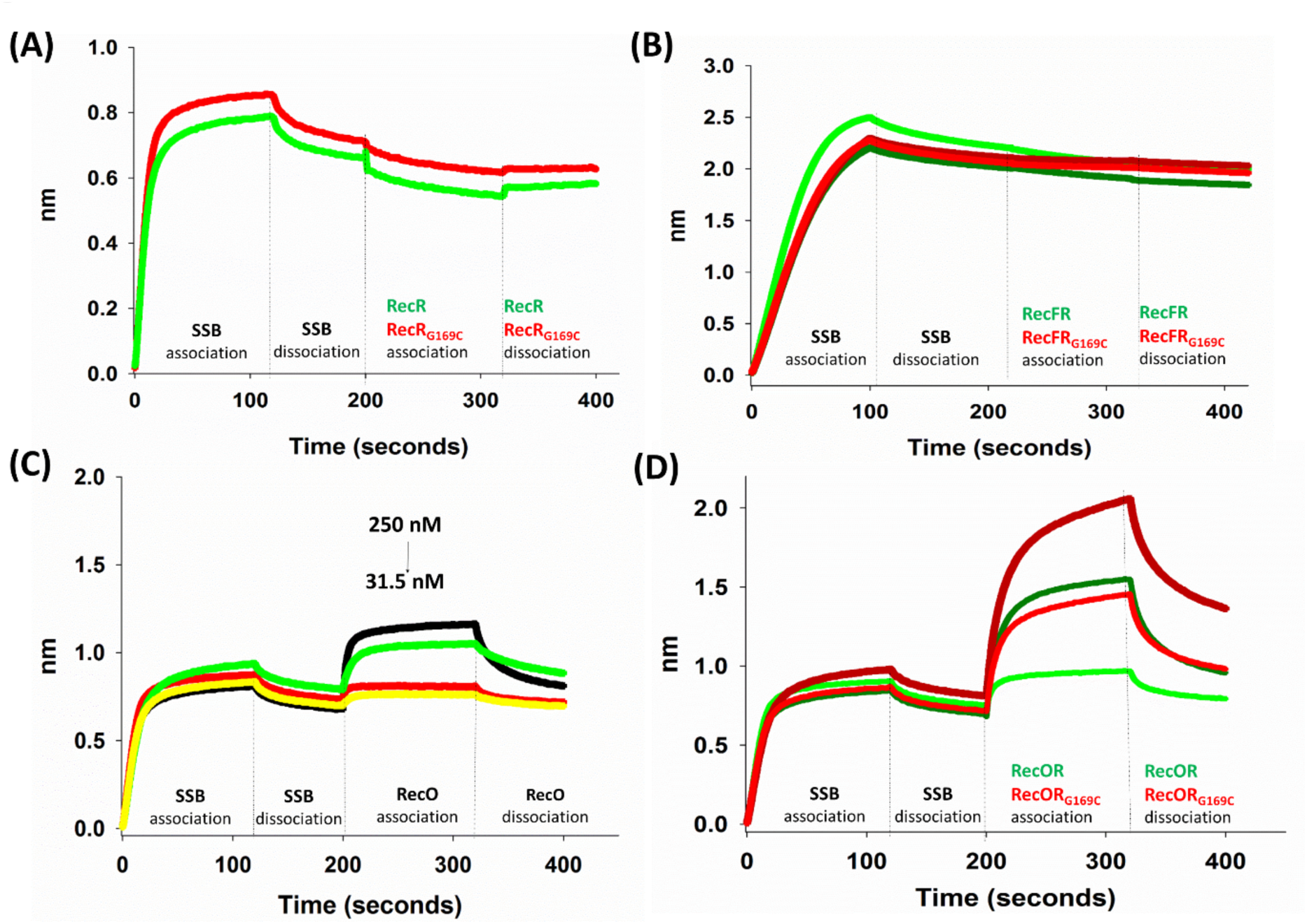
Interactions of SSB coated ssDNA with the RecFOR proteins. **(A)** Interactions of the wild type RecR (green) and mutant RecR_G169C_ (red) with the SSB coated ssDNA. **(B)** Interactions of SSB coated ssDNA with TtRecF in the presence of the wild type TtRecR and mutant TtRecR_G169C_. The dark green curve denotes incubation of the SSB coated ssDNA with 500 nM of TtRecR and 250 nM TtRecF. The light green curve denotes incubation of the SSB coated ssDNA with 250 nM of TtRecR and 250 nM TtRecF. The dark red curve denotes incubation of the SSB coated ssDNA with 500 nM of TtRecR_G169C_ and 250 nM TtRecF. The light red curve denotes incubation of the SSB coated ssDNA with 250 nM of TtRecR_G169C_ and 250 nM TtRecF. **(C)** Interactions of the SSB coated ssDNA with different concentrations of TtRecO are represented by colored curves. **(D)** Interactions of the SSB coated ssDNA with the TtRecO in the presence of the wild type TtRecR and mutant TtRecR_G169C_. The dark green curve denotes incubation of the SSB coated ssDNA with 500 nM of TtRecR and 250 nM TtRecO. The light green curve denotes incubation of the SSB coated ssDNA with 250 nM of TtRecR and 250 nM TtRecO. The dark red curve denotes incubation of the SSB coated ssDNA with 500 nM of TtRecR_G169C_ and 250 nM TtRecO. The light red curve denotes incubation of the SSB coated ssDNA with 250 nM of TtRecR_G169C_ and 250 nM TtRecO.

## Conclusion

Condition dependent stabilization of homomeric or heteromeric complexes plays a major role in regulating many major pathways in the cells. However, these complexes are very transient, which makes it difficult to study their function. Hence, one way to stabilize is through the di-sulphide bonds. RecO and RecF share a common interaction site on the RecR protein and form a hetero-hexameric complexes in a ratio 2RecO/RecF:4RecR. TtRecR exists as a dimer in solution; conversely, the crystal structure supports a tetrameric arrangement of TtRecR. The tetrameric state of TtRecR was stabilized by mutating the glycine residue at the domain swap region of the C-terminal. This resulted in both the dimeric as well as tetrameric populations in the solution. This points out that the TtRecR exist in equilibrium between dimer and tetramer in solution, however as seen in the crystal structure, with the increase in concentration, the equilibrium shifts towards the tetrameric state. In comparison to the wild type TtRecR, the mutant tetrameric TtRecR showed affinity towards the ssDNA, suggesting a different mode of interaction.

TtRecF exists as a monomer in solution, it dimerizes in the presence of ATP and shows DNA dependent ATP hydrolysis. Cysteine mutations were introduced in the two interface residues of the modeled dimeric TtRecF to stabilize its dimeric state. Surprisingly, the TtRecF cysteine mutants formed a non-canonical dimer during their purification. Interestingly, in contrast to the wild type TtRecF, the non-canonical dimeric TtRecF mutants showed affinity towards ssDNA and a concentration dependent affinity change was observed in the presence of wild type TtRecR. Interestingly, wild type RecF showed no affinity towards ssDNA. Thus, suggesting a distinct interaction mode of the TtRecF protein with the ssDNA and TtRecR protein.

TtSSB and TtRecO proteins showed affinity towards the ssDNA, nonetheless, TtSSB has hundred-fold more binding affinity than that of TtRecO. The TtSSB bound ssDNA exhibited negligible dissociation and hence is used as a template to study the binding of other cognate proteins. Individually, the wild type TtRecR and the tetrameric TtRecR or in complex with the TtRecF has no affinity for the SSB coated ssDNA. On the other hand, TtRecO binds with almost equal affinity with the TtSSB coated ssDNA as observed with the ssDNA. A higher nm shift is recorded in the presence of TtRecR/TtRecR_G169C_ by forming TtRecOR/TtRecOR_G169C_ complexes. This shift is more prominent in the case of the TtRecOR_G169C_ suggesting the importance for the tetrameric state of TtRecR in stabilizing hetero-hexameric complex and also its binding to SSB coated ssDNA.

Further, a model for SSB dissociation from ssDNA can be proposed based on our studies on protein-protein and protein-DNA interactions. The TtSSB exists as a dimer in solution and binds to the ssDNA using four OB fold domains. The TtRecO protein interacts with the C-terminal of TtSSB and has a single OB fold. The dimeric TtSSB can bind at least two TtRecO proteins and the presence of two OB folds may reduce the affinity of TtSSB for the ssDNA. This phenomenon is further reinforced by the presence of TtRecR forming the TtRecOR complex. However, our experiments suggest that the reduced affinity is not sufficient for the release of TtSSB protein from ssDNA. Nevertheless, it would be sufficient for the recruitment of the RecA protein, which may replace the bound SSB and RecOR complex by forming a nucleoprotein filament on the ssDNA.

## ACKNOWLEDGEMENTS

The authors thank Department of Science and Technology (DST) for funding. The authors thank the Department of Biotechnology (DBT), Govt. of India and the facilities offered by the center of excellence in structural biology and bio-computing, Supercomputer Education and Research Center and Department of Computational and Data Sciences, Indian Institute of Science, Bangalore. The authors thank ESRF BM-14 beam line for data collection.

## Supplementary Information

**Figure S1.**
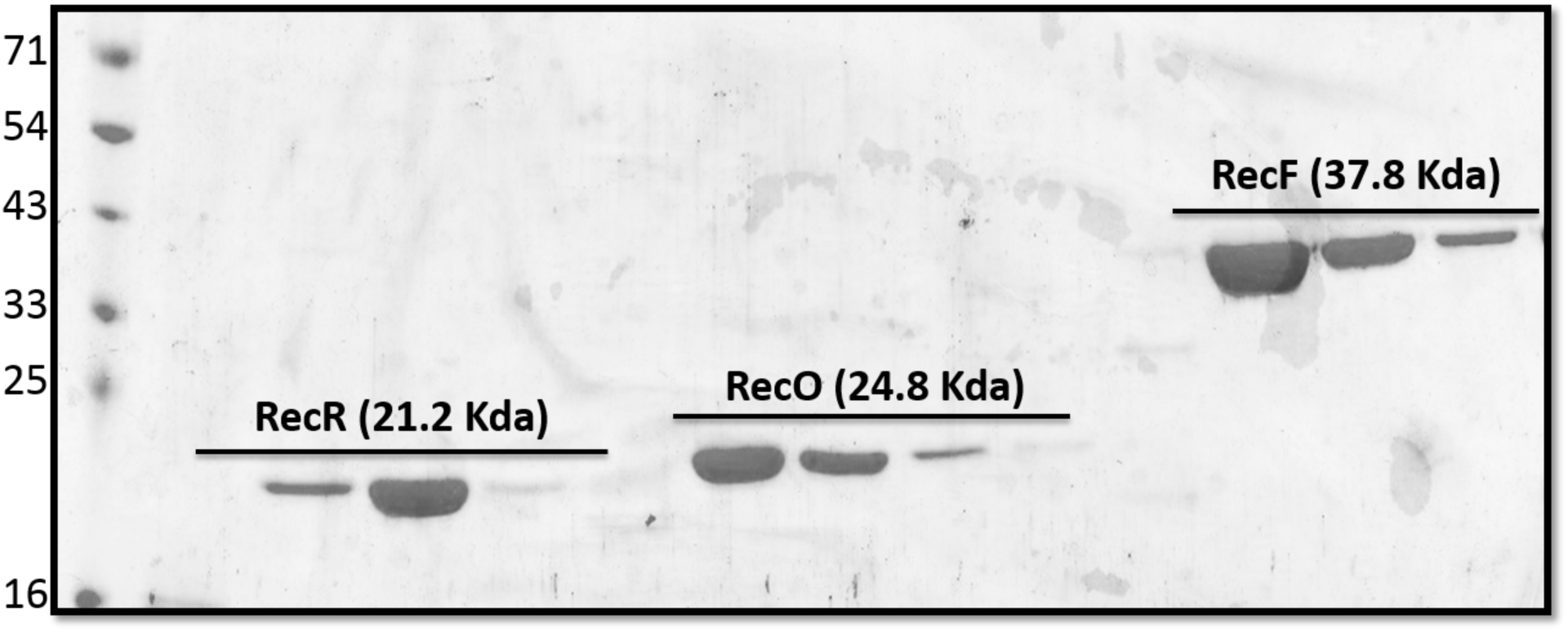
SDS-PAGE analysis of the purified TtRecR, TtRecO and TtRecF proteins.

**Figure S2.**
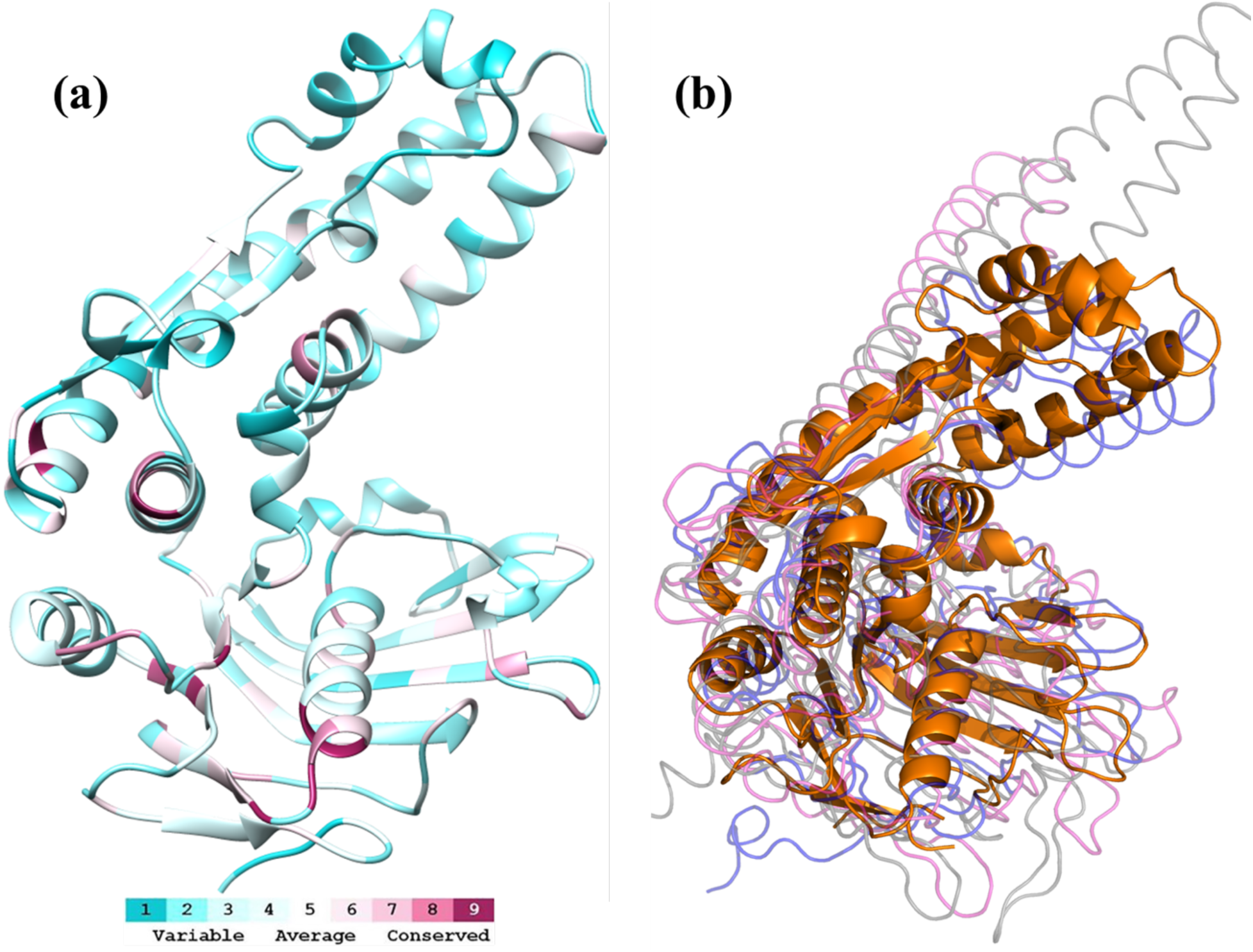
Sequence conservation and superposition of ttRecF protein with Rad50 and SMC proteins. (a) The ttRecF structure is colored based on the conservation score obtained from the alignment of ttRecF with drRecF, Rad50 (*Pyrococcus furiosus*) and SMC proteins (*Pyrococcus furiosus*). (b) Superposition of TtRecF with DrRecF, Rad50 (*Pyrococcus furiosus*) and SMC proteins (*Pyrococcus furiosus*).

**Figure S3.**
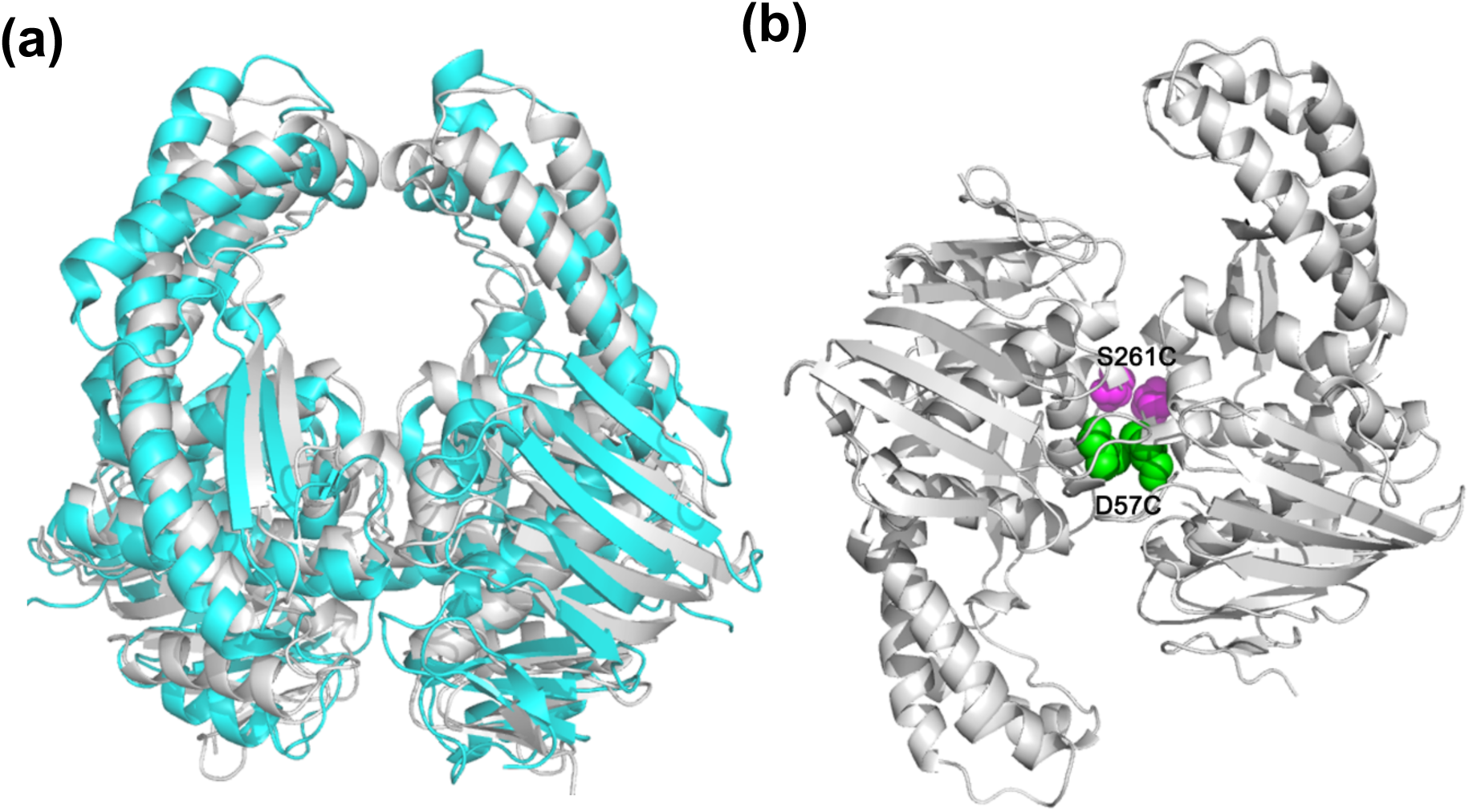
(a) Alignment of the dimeric TteRecF with the modelled dimeric TtRecF (b) The possible orientation of the mutant dimeric TtRecF_S261C_/TtRecF_D57C_ proteins.

**Table S1.**
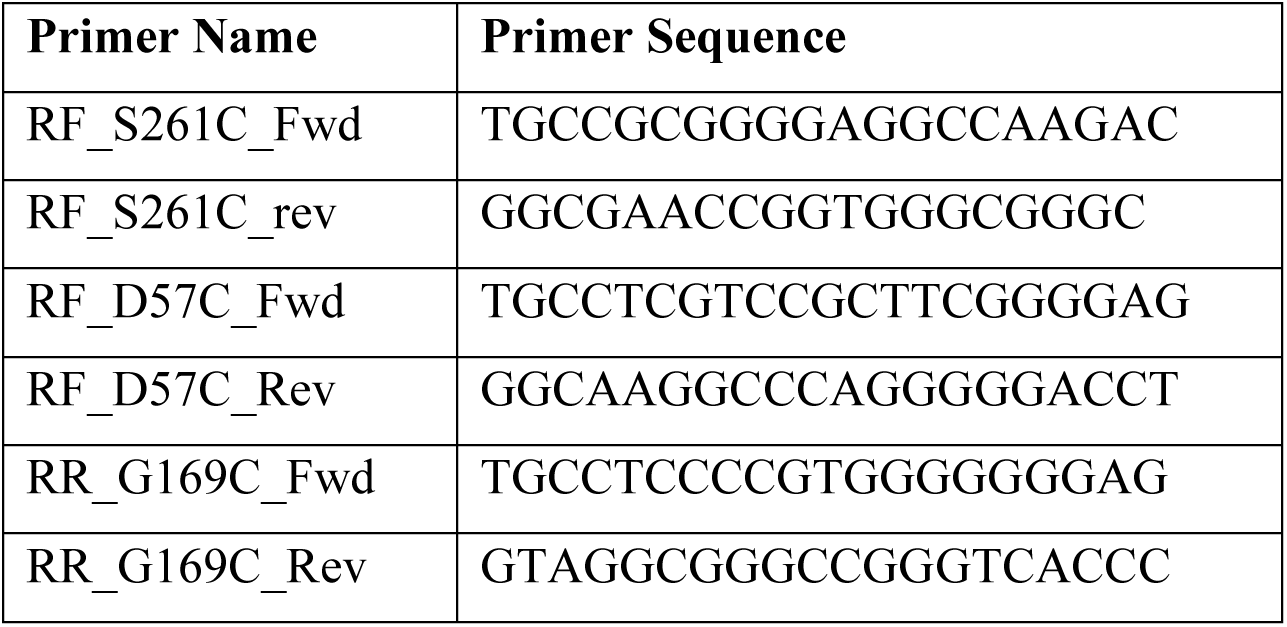
Primer sequences used for the mutagenesis in TtRecF and TtRecR proteins.

**Table S2.** Interface comparison between the dimeric TteRecF and the modelled dimeric TtRecF.

| **Title** | **TteRecF** | **TtRecF** |
| --- | --- | --- |
| Buried area upon the complex formation (Å) | 1347.0 | 2310.5 |
| Buried area upon the complex formation (%) | 3.84 | 6.77 |
| Interface area (Å) | 673.5 | 1155.25 |
| Interface area Chain A (%) | 3.83 | 6.78 |
| Interface area Chain B (%) | 3.85 | 6.77 |
| POLAR Buried area upon the complex formation (Å) | 786.0 | 1265.5 |
| POLAR Buried area upon the complex formation (%) | 58.35 | 54.77 |
| POLAR Interface area (Å) | 393.0 | 632.75 |
| NO POLAR Buried area upon the complex formation (Å<sup>2</sup>) | 561.0 | 1045.1 |
| NO POLAR Buried area upon the complex formation (%) | 41.65 | 45.23 |
| NO POLAR Interface area (Å) | 280.5 | 522.55 |
| # residues at the interface | 49 | 78 |
| Residues at the interface of Chain A | 25 | 38 |
| Residues at the interface of Chain B | 24 | 40 |

